# Three-dimensional reconstruction of fetal rhesus macaque kidneys at single-cell resolution reveals complex inter-relation of structures

**DOI:** 10.1101/2023.12.07.570622

**Authors:** Lucie Dequiedt, André Forjaz, Jamie O. Lo, Owen McCarty, Pei-Hsun. Wu, Avi Rosenberg, Denis Wirtz, Ashley Kiemen

## Abstract

Kidneys are among the most structurally complex organs in the body. Their architecture is critical to ensure proper function and is often impacted by diseases such as diabetes and hypertension. Understanding the spatial interplay between the different structures of the nephron and renal vasculature is crucial. Recent efforts have demonstrated the value of three-dimensional (3D) imaging in revealing new insights into the various components of the kidney; however, these studies used antibodies or autofluorescence to detect structures and so were limited in their ability to compare the many subtle structures of the kidney at once. Here, through 3D reconstruction of fetal rhesus macaque kidneys at cellular resolution, we demonstrate the power of deep learning in exhaustively labelling seventeen microstructures of the kidney. Using these tissue maps, we interrogate the spatial distribution and spatial correlation of the glomeruli, renal arteries, and the nephron. This work demonstrates the power of deep learning applied to 3D tissue images to improve our ability to compare many microanatomical structures at once, paving the way for further works investigating renal pathologies.

## Introduction

The kidneys, responsible for blood filtration and urine formation, are one of the most structurally intricate organs in the body. ^1^ The functional unit of the kidney, the nephron, is a convoluted and three-dimensional (3D) tube which can be sub-divided into distinct compartments: the glomerular tuft and its associated Bowman’s capsule, the proximal tubule, the loop of Henle, the distal tubules, and the collecting ducts. Each of these structures are intricately organized in relation to each other and to the renal vasculature.

The 3D arrangement and morphology of nephrons are key markers of overall renal health, and alterations to the nephron organization are routinely used to diagnose renal pathologies. For example, the Banff classification system has developed specific guidelines for the quality assessment of kidney transplant biopsies through the counting of glomeruli.^2,3^ Similarly, evaluation of chronic renal damage typically involves visually estimating the degree of fibrosis and the proportion of atrophic tubules. ^4^ Yet, our understanding of the complex organization of the nephrons is incomplete. For example, although the development of diabetic kidney disease has been associated with different markers of tubular injury and inflammation, its exact impact on the overall architecture of the kidney remains poorly understood. ^5^

Past efforts in organ mapping have been used to study kidney diseases and have generally taken one of two approaches: label many structural components in small tissue samples ^6^ or label few components in large tissue samples. ^7–16^ While these initiatives have advanced our understanding of renal diseases, we propose that the structural complexity of the kidney calls for a hybrid technique: reconstruction of large samples and microanatomical labelling of many tissue components at single-cell resolution. Recent works have demonstrated the value of 3D mapping of large biospecimens for understanding tissue structure, capturing the inter- and intra-sample heterogeneity, identifying rare events, and improving pathological classification and tumor grading. ^17–22^

In this paper, we use CODA ^23^, a novel imaging workflow that creates volumetric reconstruction of mm^3^ to cm^3^-scale tissue samples at single-cell resolution. CODA utilizes histological image registration and semantic segmentation to create 3D quantifiable maps of normal and diseased microanatomy from hematoxylin and eosin (H&E) staining alone. CODA has been applied to normal and cancer-containing samples spanning pancreas, liver, lung and skin. Here, we utilize CODA to deeply profile the microanatomy of the kidney. We extend CODA through demonstration of its ability to create whole-organ models with 17 labelled anatomical components through quantitative mapping of fetal rhesus macaque kidneys. As macaque anatomy more closely resembles human than mouse anatomy, macaques have become vital, translational model systems for the study of a wide array of human diseases including diabetes and those associated with age. ^24–26^ Additionally, a rhesus macaque model allowed histological sectioning and imaging of full organs. H&E images are the gold standard in histology.^27^ As such, we hypothesized that deep learning models could identify physiologically relevant microstructures of the kidney from H&E alone. ^28^ By doing so, we were able to expand the range of labels beyond what has been achieved with fluorescent antibodies and tissue clearing to distinguish structures such as developing and mature glomeruli. ^29^

With these reconstructed kidneys, we first deconstruct the size, cellularity, and cell density of the global and zonal regions of the kidneys. Next, we explore the relationship of glomerular size, cellularity, and developmental state to its position within the renal architecture. Lastly, by comparing our glomerular and vasculature labels, we quantify the complex relationship between the corpuscles and renal arterioles. The results of the different quantifications performed suggest a great heterogeneity in the spatial organization of developing kidneys and demonstrate the power of CODA to deeply interrogate the functional units of organs.

## Results

### 3D reconstruction of fetal kidneys at cellular resolution using CODA

The CODA imaging workflow was used to create 3D reconstructions of 4 bisected, Rhesus macaque kidney samples, collected from an 80-day old fetus (equivalent to a mid-trimester human fetus)**(Fig. 1A)**. First, a nonlinear image registration was used to align the 4μm-thick, serially cut, H&E-stained histological images into a semi-continuous stack **(Fig. 1B)**. A cell detection workflow was optimized to detect the coordinates of cell nuclei on the H&E images with a true positive rate of 84.9 %, a false positive rate of 12.4% and a false negative rate of 15%, on par with accepted validation metrics in the field (**Fig. 1C)**. ^23,30–32^ Finally, a deep learning algorithm was trained to recognize 17 microanatomical components of the developing macaque kidney **(Fig. 1D)**. Detected components included 4 developed components of the nephron (proximal tubule, loop of Henle, distal tubule, collecting duct), 3 subtypes of the vasculature (arteries, arterioles, and non-arterial vessels [veins and lymphatics]), 3 glomerular structures (mature glomerular tufts, developing corpuscles, and Bowman’s capsule & urinary space), and 2 components of the developing kidney (developing nephrons and undifferentiated blastema cells). The model, whose performances were tested on annotations unseen during the training, reached an average per-class recall of 89.7% and an average per-class precision of 90.6%. While several groups have shown success using tissue clearing or micro-CT in labelling components such as the glomerular capillary tufts,^10^ the renal vasculature,^14^ and the branching ureteric,^13^ here we were able to label all components together in a single kidney using deep learning.

**Figure 1:**
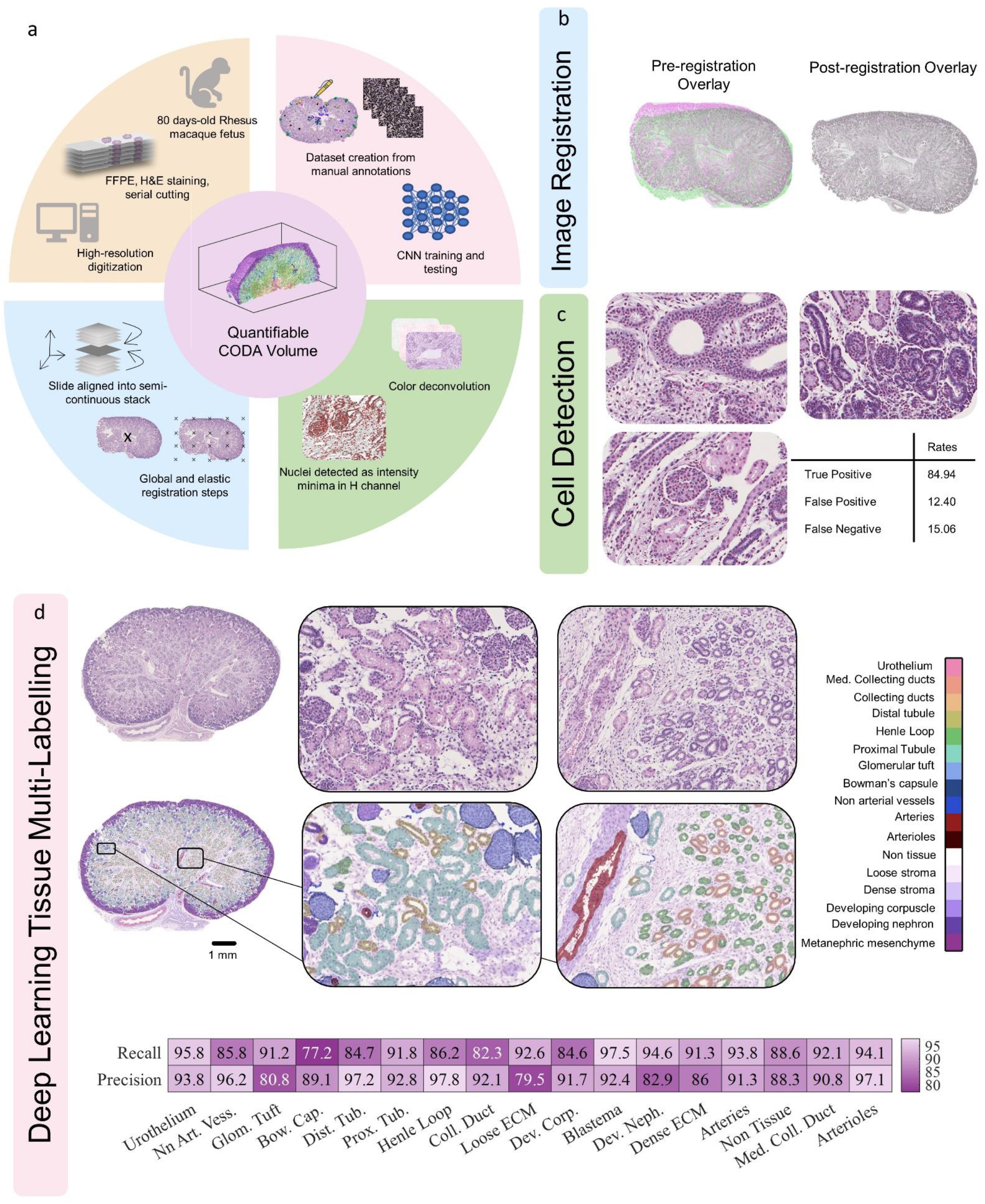
CODA. **A**. Quantifiable 3D reconstructions of FFPE, serially sectioned and H&E-stained images of fetal rhesus macaque kidneys are created using the CODA workflow. **B**. Serial H&E images were aligned using nonlinear image registration. **C**. Cell nuclei positions were determined on the hematoxylin channel of the H&E images, obtained using color deconvolution. **D**. A deep learning model was trained from manual annotations to label 17 different renal tissue subtypes on the H&E images with high accuracy.

### Gross Microanatomy of fetal rhesus macaque kidneys

The results of each CODA step were integrated to create fully visualizable and quantifiable 3D maps of the kidneys, whose volumes ranged from 19.9 to 29.7 mm^3^ and comprised between 12.8 and 17.3 million cells. (**Fig 2A**) This includes all cell nuclei detected on the H&E slide, regardless of their functional type. The color-coded rendering of one of the labeled volumes illustrates the complex and dense architecture of developing kidneys in the rhesus macaque fetus. **(Fig. 2B)** We subsequently assessed the kidney composition through quantification of the volume fraction, cellular composition (cell fraction) and cell packing density of each microanatomical structure. **(Fig. 2C)** We observed that, while the overall tissue was mostly composed of loose and dense stroma, specific microstructures, such as the proximal tubules and the metanephric mesenchyme make up the majority of the functional parenchyma (as seen in the cell fraction).

**Figure 2:**
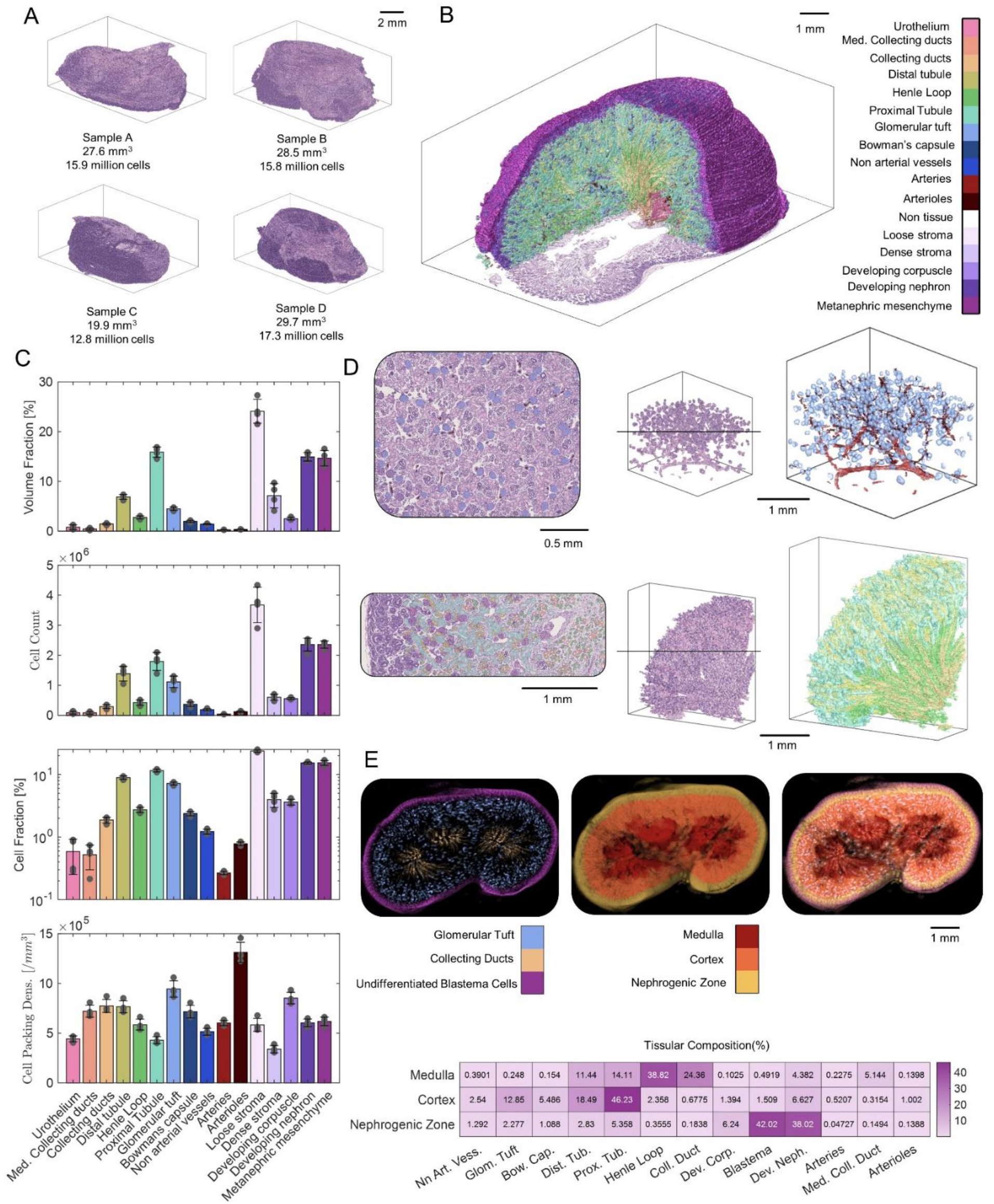
Gross microanatomy of fetal Rhesus macaque kidneys. **A**. Four half fetal Rhesus macaque kidneys were reconstructed, and their volumes and total cell count were assessed. **B** The 3D reconstructions include 17 microanatomical labels. **C**. The volume fraction, cell count, cell fraction and cell packing density of each microanatomical subtype were computed and compared. **D**. A 3D rendering of glomerular tufts with arteries and arterioles and a 3D rendering of the different parts of the tubules were compared to their corresponding aligned H&E volumes. Examples of classified H&E images with segmented structures are shown for validation. **E**. The three zones of the developing kidney were delineated based on the presence of three defining elements. The location of the zones and their corresponding defining structures were visualized using z-projections. The tissular composition of each zone was assessed.

Physiologically relevant combinations of microstructures were then isolated from the general volume and rendered in 3D. First, we rendered the glomerular tufts, arteries, and arterioles to visually demonstrate the spatial correlation between these functional structures of the kidney. Second, we rendered the four detected sections of the renal tubule (collecting ducts, distal tubule, Henle loop and proximal tubule) to show the very specific co-localization of these structures in the kidney, distinctly highlighting the cortex and the medulla. The equivalent three-dimensional H&E volume was created for both combinations of structures and rendered as a comparison, along with 2D H&E images where each of these structures were selected to additionally showcase the exquisite detail of the deep learning segmentation. **(Fig. 2D)**

The three distinct anatomical regions of the fetal macaque kidney are the nephrogenic zone, the medulla and the cortex. We quantitatively defined each zone in 3D space based on the presence of undifferentiated blastema cells, glomerular tufts and collecting ducts. respectively. **(Fig. 2E)** Assessment of the tissular composition of each zone showed that the medulla was mostly composed of Henle loops and collecting ducts, the cortex was mostly composed of proximal tubules, and the nephrogenic zone was mostly made of undifferentiated blastema cells and developing nephrons. Large arteries were mostly found in the cortex and the medulla while smaller arterioles were mainly found in the cortex. The cortex contained the highest fraction of non-arterial vessels (veins and lymphatic vessels). Interestingly, the medulla is the least vascularized zone, due to its very low levels of non-arterial vessels in comparison to the other two zones.

### Spatial arrangement of developing and mature glomeruli in fetal macaque kidneys

We then moved our analysis beyond the global and regional quantifications to assess the spatial organization of different structures relevant to kidney function. We started by considering glomeruli and their spatial distribution in the kidney during crucial developmental phases. Glomeruli are a widely accepted marker of renal health ^2,33^and have been extensively investigated. ^9–11,34^Aside from the number and size of glomeruli on a whole organ level, which is already challenging information to extract using most organ mapping techniques, we used CODA to assess parameters that are difficult to quantify at scale using conventional techniques. These parameters included the volume occupied by the Bowman’s capsule and the urinary space in each glomerulus, as well as the absolute number of cells and the cell packing density of glomeruli across different regions of the kidney.

The thickness of the combined Bowman’s capsule and urinary space (referred to as the Bowman’s capsule thickness in the remainder of this paper) was assessed in every detected mature glomerulus. **(Fig. 3A)** The Bowman’s capsule thickness displayed a relatively spread-out distribution, with a minimum value of 0 μm and maximum value of 6 μm, but with a mean value of 2 μm. This parameter was evaluated against others in each glomerulus, and it appeared that; larger glomeruli (by volume) had a larger Bowman’s capsule thickness and glomeruli located further from the surface of the kidney had a larger Bowman’s capsule thickness. No relationship was found between Bowman’s capsule thickness and glomerular cell packing density.

**Figure 3:**
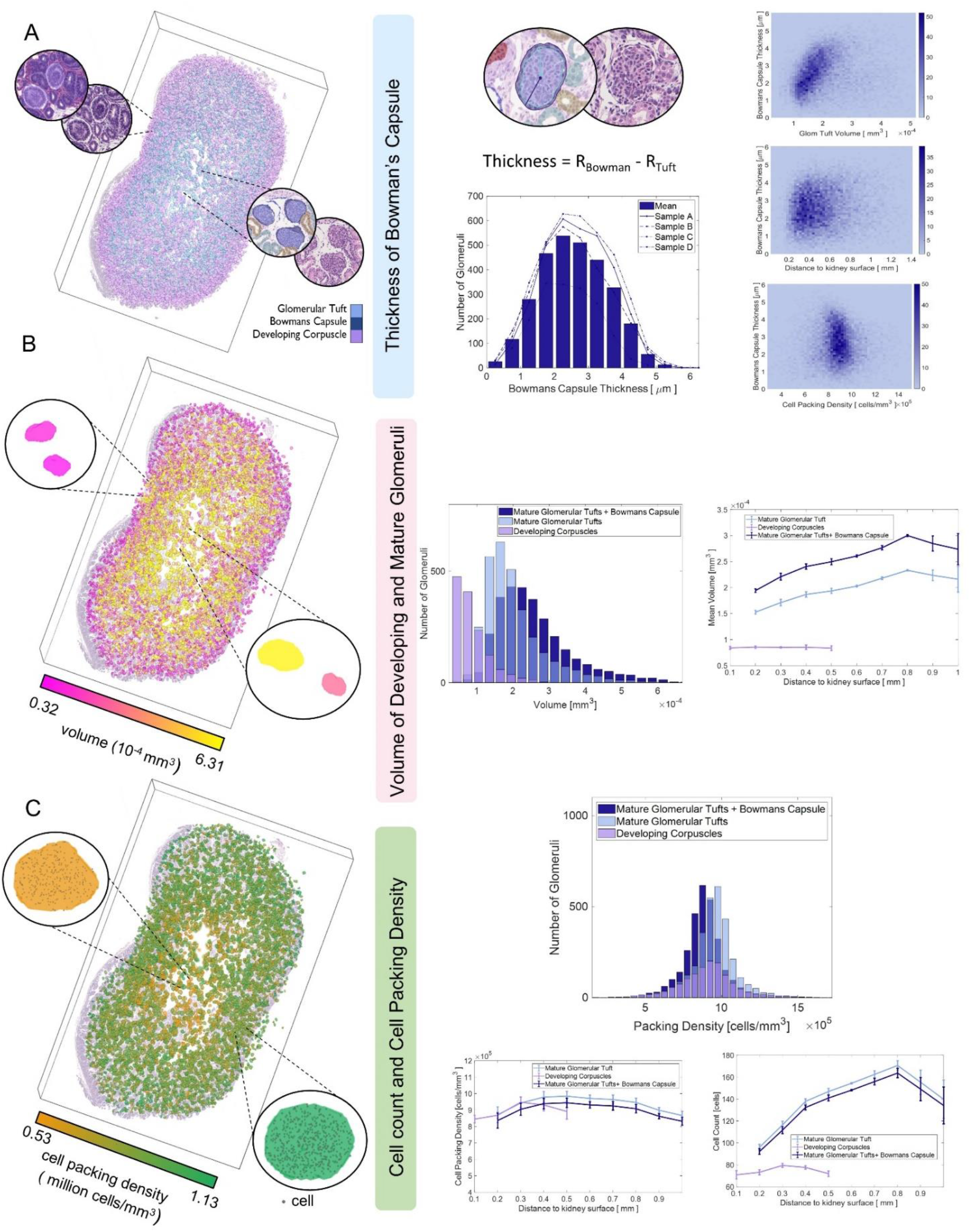
Spatial distribution of developing and mature glomerular tufts in the fetal Rhesus macaque kidney. **A**. The thickness of the Bowmans’ capsule and the urinary space was assessed in mature renal corpuscles. The distribution of thickness was computed in the four samples and the thickness was evaluated as a function of different parameters. **B**. A three-dimensional heatmap of glomerular volumes was rendered. The distribution of volumes in mature and developing glomeruli was described and the relationship between the volume of a corpuscle and its position in the organ was evaluated. **C**. A three-dimensional heatmap of glomerular cell packing densities was created. The distribution of cell packing densities for the three glomeruli populations was computed. The cell packing density and absolute cell count of each glomerulus were studied as functions of the glomeruli’s distance to the surface of the kidney.

The volume distribution of developing corpuscles and mature glomerular, with and without their respective Bowman’s capsule and urinary space, was assessed. **(Fig. 3B)** As expected ^34^, developing corpuscles displayed much smaller volumes than mature glomeruli. It appeared that the distribution of mature glomerular tufts was relatively narrow with a mean volume of around 1.75x10^5^ μm^3^. Interestingly, consideration of the combined volume of the glomeruli and the associated Bowman’s capsule thickness broadened the range of the distribution, suggesting that glomeruli of similar volumes possess a range of Bowman’s space. From the 3D volume heatmap with all the glomeruli, in which each glomerulus was assigned a color based on its size, we note that most of the smaller corpuscles lie on the margins of the kidney, while larger glomeruli localize nearer the center of the kidney. This observation conforms to previous assessments of the spatial distribution of glomerular size. ^11^Additionally, as we can distinguish developing and mature glomeruli, we note that developing corpuscles primarily occupy space within 0 to 0.5 mm of the kidney surface, and that, during the developmental stages, the size of the corpuscle is not related to its location relative to the surface of kidney. In contrast, mature corpuscles seem to increase in size as a function of their distance to the kidney surface, irrespective of consideration of the Bowman’s capsule thickness.

We next characterized renal corpuscles in terms of cell packing density. **(Fig. 3C)** Similarly to Fig 3B, we described the distributions of cell packing density of developing and mature corpuscles, with and without considering the Bowman’s space. Distribution of cell packing densities of the three glomeruli populations were relatively similar. It also appeared that average cell packing density of a glomerulus does not vary as a function of its distance from the surface of the kidney, although it is visible from the 3D cell packing density heatmap that the glomeruli with the lowest cell packing density were predominantly located deep in the cortex. Interestingly, while there is almost a 20-fold increase between the lowest and highest glomerular volumes detected, the highest and lowest cell packing densities varied by only 2-fold. We conclude that while the size of developing and mature glomeruli varies greatly, the cell number varies linearly with this size, such that cell density remains consistent. While developing corpuscles have a much lower absolute cell count than mature ones, they also have a much smaller volume.

### Characterization of the 3D organization of renal arterial vessels and their association with glomeruli

The network of renal arterial vessels, split into arteries and arterioles, was 3D reconstructed **(Fig. 4A)**. Using the 3D arterial data, we assessed the volume fraction of arterial vessels as a function of the branching point of the segmental artery (see black dot indicator in Fit 4A). First, we noted that the fraction of large arteries was highest at the vascular entrance to the kidneys and decreased steadily as a function of distance into the kidney. In contrast, the size of arterioles remained constant across the renal architecture. We also considered the relationship of vessel diameter as a function of the distance from the branching point of the segmental artery and similarly found that arteries are largest when entering the kidney before decreasing in size, while the arterioles maintain constant diameter throughout the kidney. ^7,8^

**Figure 4:**
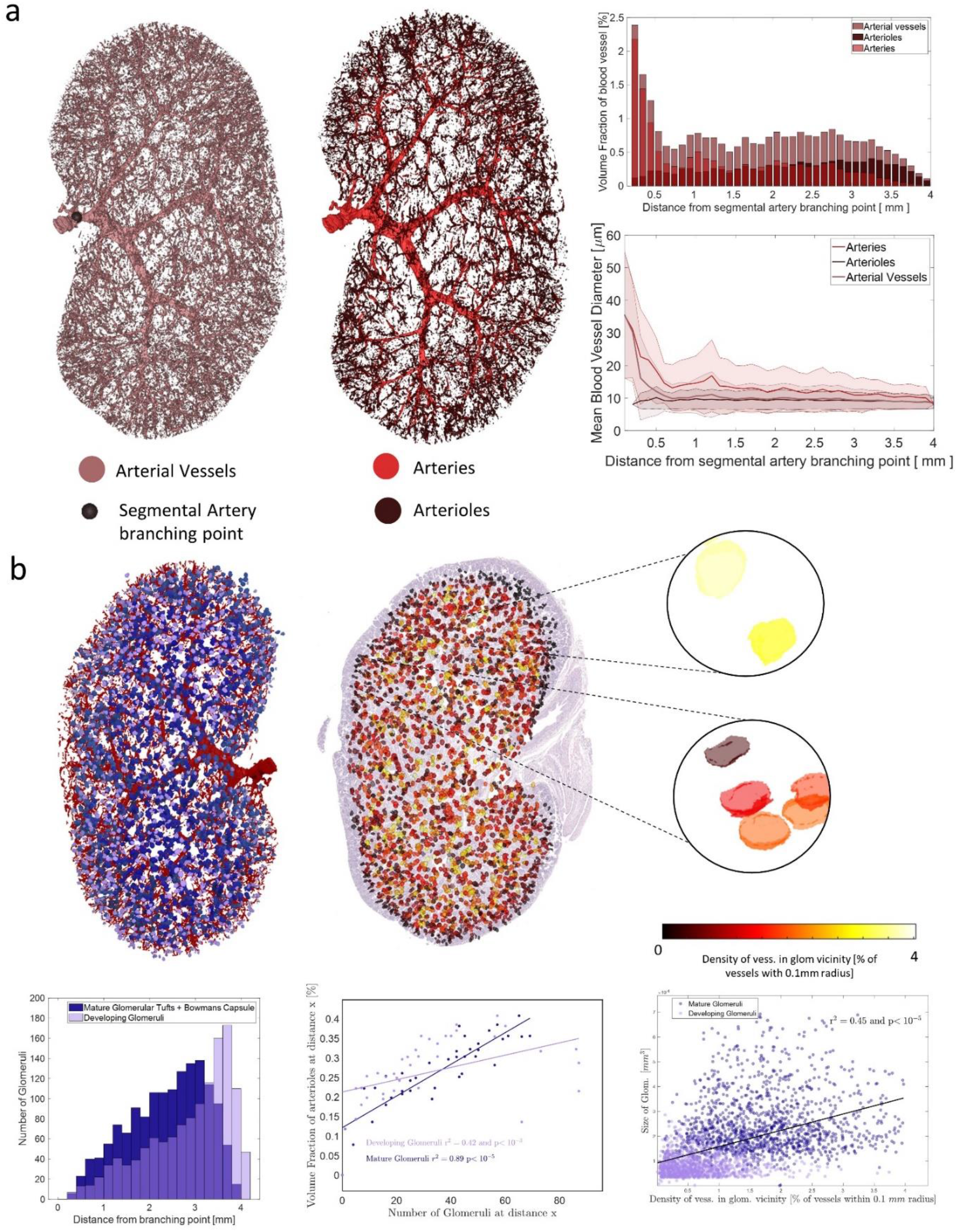
3D organization of arterial vessels in the fetal macaque kidney and their association with glomeruli. A. Arteries and arterioles were reconstructed in three dimensions. The volume fraction and the mean vessel diameter of vessels were assessed as a function of the distance from the branching point of the segmental artery. B. The association between arterial vessels and glomeruli was determined through comparison of the density of vessels in the vicinity of renal corpuscles and the distribution of glomeruli away from the arterial branching point. The number of glomeruli detected within incremental distance windows (referred to as ‘distance x’ in the axis label) was evaluated against the fraction of the volume occupied by arterioles within the same distance window (referred to as ‘distance x’ in the axis label). Glomeruli sizes were described as a function of the density of arterial vessels in their vicinity.

Finally, we combined our analysis of the renal arterial vessels with our assessment of glomeruli. **(Fig. 4B)** We investigated the distribution of mature and developing corpuscles as a function of their distance from the branching point of the segmental artery. In addition, we defined what we term “glomerular vascularization” as the volume fraction of arterial vessels within a 0.1 mm radius around each point of a glomeruli. We found a strong correlation between glomerular size and degree of vascularization (r^2^ = 0.45 and p< 10^-5^), suggesting that larger glomeruli are more vascularized than smaller glomeruli. Next, we compared the glomerular density to the volume fraction of arterioles at different distances from the vascular entrance to the kidney to reveal that the number of (developing and mature) glomeruli at different regions in the kidney is highly correlated with the volumes of arterioles in the same regions (r^2^ = 0.76 and p< 10^-5^). This same analysis revealed that developing glomeruli were less vascularized than mature glomeruli, emphasizing the importance of differentiating developing and mature structures in study of fetal tissues.

## Discussion

In this work we created cellular-resolution maps of fetal rhesus macaque kidneys to explore the inter-relation of renal vascular, glomerular, and nephron structure. Through successfully labelling seventeen microanatomical structures in renal histology at >90% accuracy, we demonstrate the marked power of deep learning in creating highly annotated 3D tissue maps for quantitative interrogation of tissue architecture. Importantly, we demonstrate that deep learning can aid in differentiation of structures that cannot be distinguished using conventional tissue mapping techniques such as molecular staining. Using histological morphology to guide a deep learning algorithm, we differentiate large arteries, arterioles, and non-arterial vessels. We additionally differentiated developing vs. mature glomeruli, and we identified Bowman’s space: an empty cavity between the Bowman’s capsule and the glomerular tuft that, by definition, cannot be molecularly labelled.

This advancement is important in enabling quantitative comparison of functionally important but molecularly indistinguishable structures, as demonstrated though our comparison of the spatial position within the kidney of developing vs. mature glomeruli and arterial vessels.

By computing the respective tissular compositions of the nephrogenic zone, the cortex, and the medulla, we show the differing composition of microanatomical structures across the zones of the kidney. The developing nephrons and undifferentiated blastema cells were majority components in the nephrogenic zone, while deeper in the kidney, mature components such as glomeruli and proximal were more prevalent in the cortex and loops of Henle and collecting ducts were more prevalent in the medulla. We also observed differential vascularization of the different anatomical regions of the developing kidney through the computation of their respective volume fractions in arteries, arterioles, and non-arterial vessels. Examining the association between mature and developing glomeruli and arterial vessels in a spatially resolved manner allowed us to show that developing glomeruli were less vascularized than mature ones. While extensive work has not been performed that could validate our findings concerning developing structures, our observations of vascular distribution in the kidney were on par with previous work showing that blood vessels do not associate uniformly with different other renal structures. ^35^

Overall, we illustrate the value of deep learning-based tissue mapping through construction of single-cell resolution 3D maps of kidneys with numerous microanatomical structures labelled. We revealed the marked spatial organization of developing and developed renal structures, with the developing regions generally nearer the outer margins and the mature components at the core. Together, this work demonstrates the power of CODA in mapping numerous, inter-related anatomical components in complex organs such as the kidney.

## Authors Contribution

A.L.K. and D.W. conceived the project. J.L. and O.M. provided the biospecimens. L.D. led the histological image processing, visualizations, and representation of results, with assistance from A.F. L.D. created the deep learning model of renal architecture, with extensive pathology guidance from A.R. L.D. wrote the first draft of the manuscript, which all authors edited and approved.

## Acknowledgements

The authors acknowledge the following sources of support: OD011092; U54CA268083;U54AR081774; Lustgarten Foundation-AACR Career development award for pancreatic cancer research in honor of Ruth Bader Ginsburg; Susan Wojcicki and Denis Troper.

## Materials and Methods

### Sample Acquisition

Kidneys were collected from a male rhesus macaque fetus of 80 days gestation (equivalent to a mid-second trimester human fetus) at the Oregon National Primate Research Center (ONPRC). The collected samples were then fixed in formalin, embedded in paraffin and sectioned every 4μm. The resulting slides were subsequently stained using H&E and scanned at 20× using a Hamamatsu NanoZoomer.

### Image Registration

Openslide software was used on the original images scanned at 20× (corresponding to 0.5 μm/pixel) to create down-sampled versions of each image, with a resolution of 8um/pixel, via nearest neighbor interpolation.

The down-sampled images were then registered with a combination of global and elastic registration using the method described in ^23^.

Briefly, the down-sampled grayscale and gaussian-filtered versions of the high-resolution images were registered using global and elastic registration with the center image of the stack as a point of reference. The parameters for global registration, the rotation angle and the translation, were sequentially computed for an image and the next three images closest to the center image. The rotation angle was determined first by selecting the angle between 0 and 359 that lead to the highest cross-correlation between Radon transforms of the images being registered and the translation was then found by maximizing the cross-correlation between rotated images. The final global registration was chosen as the one yielding the highest pixel-to-pixel correlation between moving and reference images among the three that were computed, while the other two were discarded. This was used to ensure no compounding error would arise from potential folding or splitting of tissue regions on certain slides. Elastic registration was then subsequently computed on the globally registered images by dividing them into smaller tiles on which global registration was performed. Interpolation and Gaussian-smoothing of these local global registration results yielded the nonlinear, elastic registration transformation. Successive global and elastic registration were computed for all images of the samples, such that they would be aligned in the same coordinate system.

### Nuclei cell detection on histological images

Openslide software was used on the 20× images to downsample them to 2μm/pixel images, which were used to determine the position of cell nuclei by following the method introduced in ^23^First, color deconvolution was applied on the colored H&E images to extract the hematoxylin channel. To do so, the H&E2 built-in staining matrix of the ImageJ Color Deconvolution 2 plugin was used to deconvolve the H&E images into their eosin, hematoxylin, and background channels. Following the method described in ^36^, the hematoxylin channel images were smoothed, and the position of each cell nuclei was identified as the 2D intensity minima of the resulting images. The true positive, false positive and false negative rates of the detection were computed by comparing the automatically generated coordinates with manually annotated ones on 16 tiles 4mm2, created from an annotation function built in Matlab 2021b. The maximum distance between an automatically generated set of coordinates and the annotated ground truth for it to be considered a true positive was set to be 7μm.

### Tissue labeling using a deep learning semantic segmentation model

Following the same method as in ^23^, a deep learning semantic segmentation model was trained and validated to recognize different microanatomical renal structures on reduced size copies of the original 20× images, corresponding to 2μm/pixel.

Thirteen 20× images equally spaced in the stack were used for each sample. These images were annotated using Aperio ImageScope with between 30 to 100 examples of each class of microanatomical components. Manual annotations were stored in a separate xml file containing the annotations coordinates for each annotated image. These coordinates were then overlaid on the 2μm/pixel images and were used to create the bounding box of each annotation. A matrix was created to keep track of what type of tissue each annotation contained.

Training and validation images were created from 9000×9000×3 zero-value RGB tiles which were iteratively recovered with annotation bounding boxes, until it was covered at more than 65%. Annotations were randomly selected from the bounding boxes containing the least represented class, so that all classes ultimately accounted for a similar number of pixels on the filled tile image. Some of the annotations also undergone different data augmentation operations, such as rotation, scaling and hue augmentation. This process was used to create 8 9000×9000×3, one was used for validation and the remaining 7 were used for training. Each tile was cut into smaller 144 750×750×3 images, which created a training dataset of 1,008 images and a validation dataset of 144 images.

These image sets were then used to train and validate a resnet50 network adapted for DeepLab v3+. The precision and recall of the trained model were assessed on 2 manually annotated images that were not seen during the training or the validation. New annotations were added and used to train a new model until the accuracy of each tissue class reached over 75%.

The final model was then used to semantically segment all the 2μm/pixel resolution images of the samples.

### 3D reconstruction of samples

Similarly to the way it was done in^23^, the samples were reconstructed as 3D matrices, built by applying the image registration transformations to the semantically segmented images and assembling the registered classified images in the third dimension. These 3D matrices were then rendered in MATLAB 2023a using the patch and isosurface commands. To help with the visualization, each tissue subtype was assigned a unique RGB color triplet. In parallel, a 3D cell matrix was created for each sample by registering the coordinates of the detected cell nuclei detected and consolidating the results in 3D. The 3D cell matrices were visualized as single cells using the scatter3 function in MATLAB 2023a.

The 3D reconstructions had an original resolution in *xy* of 2μm/voxel, corresponding to the resolution of the 10× images used for the cell detection and the deep learning segmentation, and a resolution in z of 4um/voxel, corresponding to the thickness of a histological section. However, the volumes were downsampled to an isotropic resolution of 4×4×4 μm^3^/voxel, using nearest neighbor interpolation, for the rest of the computations.

### Calculation of tissue volume, volume fraction, bulk cell packing density and local cell packing density

The volume of tissue in each sample was computed as the number of voxels labeled as tissue in the 3D matrices multiplied by the volume of a voxel, that is 4×4×4μm^3^. The volume fraction of a particular tissue subtype was then computed as the ratio between the number of voxels in the 3D matrix labeled as that class divided by the total number of tissue voxels in the volume.

The cell counts in each tissue subtype were computed by associating the detected cell coordinates with a tissue subtype in the labeled 3D matrix. Therefore, a cell nucleus detected at a voxel that is labeled as *proximal tubule* will be counted towards the number of cells for the *proximal tubule* class. The cell count of each tissue subtype was corrected using the formula introduced in (Kiemen et al., 2022),to avoid the potential double counting of nuclei detectable on multiple consecutive slides. This formula makes use a parameter D_subtype_ which was computed, for each tissue subtype, by averaging 10 manually measured diameters obtained through Aperio ImageScope. Similarly to the volume fraction, the cell fraction of each tissue subtype was computed as the ratio between the cell count of that subtype and the total number of cells detected in the sample. Finally, bulk cell packing densities were calculated, for each tissue class, as the ratio between its cell count and its total volume.

### Delineation of the zones in the developing kidney

A 3D matrix where each voxel was attributed one of the three zones of the developing kidney (the nephrogenic zone, the cortex or the medulla) was created. A zone was assigned to each voxel based on which of the undifferentiated blastema cells, the mature glomerular tufts or the collecting ducts class was present in majority within a 150μm radius around that voxel. If the percentage of undifferentiated blastema cells was the highest, the voxel was set to be in the nephrogenic zone while if it was the mature glomeruli or the collecting duct percentage, the zone was defined as the cortex or the medulla respectively.

Practically, this was done by defining, for each of the three tissue subtypes, a logical 3D matrix with positive voxels where that tissue was detected. Then, a 3D unit sphere of a 150μm radius was created and convoluted around that logical matrix. The 3D matrix resulting from this convolution operation was then divided by the volume of the sphere. This created a 3D matrix whose value at a particular voxel corresponded to the percentage of the 150um sphere centered at that voxel that is occupied by that particular tissue class. By repeating this for the two remaining classes, we were able to define the 3D matrix by assigning, at each voxel, the zone that corresponded to the tissue class with the highest percentage of presence.

### Construction of z projections

As it was presented in ^23^the Z-projection of a particular tissue subtype was created, at each pixel of a the xy plane, by summing in the z direction the voxels of the 3D matrix corresponding to that tissue class. The z-projections of different tissue classes were combined, normalized by their maximum and visualized, in the same color scheme as previously defined, with the images Matlab2023b command.

### Analysis of the spatial arrangement of mature and developing glomeruli

The developing glomerular tufts, mature glomerular tufts labels and the combined urinary space and bowman’s capsule label were isolated from the 3D matrices to create new volumes containing only these elements for each sample. These matrices underwent slight smoothing to remove any noise due to potential misclassifications by the deep learning model.

First, the thickness of the Bowman’s capsule and the urinary space associated with each individual glomerular tuft was assessed. This parameter, called Bowman’s capsule thickness for simplicity, was defined as the difference between the radius of the sphere formed by the combined bowman’s capsule and urinary space and the radius of the sphere formed by the developing/mature glomerular tuft. This parameter was assessed and compared against other parameters describing each individual glomerulus: the size of its tuft, its distance to the surface of the kidney and its cell packing density. The size and cell packing density of each glomerulus were computed as explained previously. The distance to the kidney surface was defined as the distance, computed using the bwdist function in MATLAB, between the centroid of the object formed by a glomerulus (determined using the regionprops3 function in MATLAB) and the outer surface of the whole tissue sample.

The size and cell packing distributions of glomeruli were computed. The size, cell count and cell packing density of each glomerulus was then also compared against its position with respect to the surface of the kidney. Three dimensional heatmaps of the glomerular volume and cell packing densities were built by assigning a color to each glomerulus based, respectively, on its individual size and cell packing density.

### 3D reconstruction and characterization of the arterial network

The arterial network was reconstructed from a second deep learning model able to recognize three different microstructures: renal tissue without arterial vessels, arterial vessels and their lumen, and the background of images. This deep learning model was trained and tested on a new set of manual annotations following the same pipeline as the previous 17-classes one. This was done to improve the segmentation accuracy of the original model, especially in the segmentation of the vessels lumen and smaller arterioles, which allowed to increase the resolution of the reconstructed arterial network. The new model achieved an average precision and recall of 95 %.

The results of the new 3-classes model were then consolidated using the image registration results previously computed. This allowed the creation of the 3D arterial network of each sample, with a resolution of 4×4×4μm^3^/voxel.

The newly reconstructed arterial vessels volume was smoothed using the morphological opening and closing functions in MATLAB. The arterial vessels detected on the renal capsule and around the ureter were manually excluded to keep only the inner network.

The vessels in that volume were given an artery/arteriole label by extrapolating the reconstructed results of the 17-classes model. This was done by isolating the artery and arteriole label from the full 3D reconstruction and applying the dilatation and morphological closing openly available on MATLAB. The dilated artery-arteriole volume was then dot multiplied with the cleaned arterial vessels volume. It was estimated that around 85% of all voxels in the arterial vessel reconstruction could be assigned a label from the old classification.

The volume fraction respectively occupied by arterial vessels, arteries and arterioles for different incremental windows of distance from the branching point of the segmental artery was computed. That branching point was manually determined by locating the branching point of the renal artery from the stack of aligned images. It was then defined as a sphere of 0.1-mm radius centered on the manually determined coordinate within the arterial vessel volume. The distance between the sphere and each voxel in the volume was determined using the bwdist function in MATLAB. The volume fraction of a given type of arterial vessel within a window was computed as the number of voxels detected as that vessel type, divided by the total number of voxels in the window.

Using the bwdist results previously computed, the mean diameter of the three types of arterial vessels was characterized also as a function of their distance to the branching point of the segmental artery. The diameter of the vessels were determined by applying the bwdist Matlab function on the opposite of the arterial network volume. The result of that operation was then multiplied by 2 and dot multiplied with the morphological skeleton of the arterial network (obtained using Matlab’s bwskel function). This gives a 3D skeleton in which each voxel gets its diameter as a value. The mean blood vessel diameter within a given window from the segmental artery branching point is then computed as the average of the value of the diameter skeleton voxels located within that window. This was done in turn for the three types of vessels.

### Analysis of the association between glomerular tufts and the arterial network

The association between glomeruli and the arterial network was then assessed. First, we counted the number of mature and developing glomeruli within incremental distance windows from the segmental artery branching point.

We also described how the number of the different types of glomeruli detected within a given distance window was associated with the volume fraction of arterioles previously computed within that window.

We then evaluated how the size of each glomerulus related to the density of arterial vessels within a 0.1mm radius around it. This was done by defining 0.1mm radius sphere object and convolving it around the arterial vessel volume. By dividing the result of that convolution by the volume of the sphere, we obtained at each voxel in the volume the volume fraction occupied by arterial vessels within a 0.1mm sphere centered at that voxel. The density of vessel for a given glomeruli was then obtained by averaging the densities computed for each voxel of that glomeruli.

We finally created a three-dimensional heatmap of the densities of vessels in the glomeruli vicinity by attributing a distinct color to each glomerulus based on their density value.

